# Improved detection and classification of plasmids from circularized and fragmented assemblies

**DOI:** 10.1101/2022.08.04.502827

**Authors:** Matías Giménez, Ignacio Ferrés, Gregorio Iraola

## Abstract

Plasmids are mobile genetic elements important for bacterial adaptation. The study of plasmids from sequencing data is challenging because short reads produce fragmented assemblies, requiring of subsequent discrimination between chromosome and plasmid sequences. Although circularized assemblies are now possible using long-read data, there is still a need to differentiate plasmids from other circular elements. Here, we present plaSquid, a dockerized tool developed in Nextflow that expands plasmid detection and improves replicon typing and mobility groups classification schemes, outperforming previously available methods in both precision and sensitivity. When applied to ∼10.5 million metagenomic contigs, plaSquid revealed a 2.7-fold increase in plasmid phylogenetic diversity. Also, we used plaSquid to uncover a significant role of plasmids in the widespread distribution of clinically-relevant antimicrobial resistance genes in the built environment, from cities to spacecraft. Together, we present an improved approach to study plasmid biology from fragmented or circularized genomic and metagenomic assemblies.

## Introduction

Plasmids are extrachromosomal genetic elements that replicate independently from the bacterial chromosome and are transferred and maintained in different host species. Although plasmid genes are considered non-essential for bacterial viability, many plasmid-encoded traits have an impact on bacterial adaptation as they confer resistance to environmental pressures, like antibiotics, or give access to new niches by providing specific metabolic capacities^1–3^.

Plasmids have a modular structure defined by different functionally-related gene clusters^4,5^. Replicons are composed by counter-transcript RNAs (ctRNAs), iterons, promoters and their encoded proteins, and regulate plasmid replication and copy number by interacting with the bacterial chromosome-encoded molecular machinery. Replicons are the only indispensable module to define a plasmid. Based on their sequence diversity, plasmids have been classified according to different replicon (REP) types which can be informative about plasmid host range, copy number and size^6,7^. The mobility module is present in some plasmids and confers the capacity of autonomous mobilization between hosts. Based on sequence similarity on the N-terminal portion of relaxase genes, mobilizable plasmids can be classified into mobility (MOB) groups^8,9^. These groups are phylogenetically coherent but in certain cases not entirely correlated with REP types^6^. The adaptation module consists of a variable set of elements including protein-encoding genes, transposable elements and integrative sequences that increase genetic fluidity between bacterial genomes driving their adaptation to environmental changes^10^.

High-throughput sequencing of bacterial isolates and metagenomic samples from diverse environments is enabling the massive exploration of plasmid genetic diversity, stressing their importance in infectious diseases as they frequently encode antimicrobial resistance (AMR) or virulence traits^3^. This has motivated the need for methods and schemes for plasmid identification and classification from sequencing data. Currently, plasmid classification based on MOB groups and REP types is the standard, but this approach fails to classify an important share of plasmids present in current databases^6^. Therefore, other schemes that are based on whole-sequence comparisons and network analysis have been developed^11,12^. However, there is still a need for improved methods for plasmid identification and classification from short-read data, since this requires discriminating between chromosome and plasmid sequences when working with fragmented genomic or metagenomic assemblies. Although more contiguous or complete assemblies are now possible using long-read sequencing technologies, there is still a need for approaches that can identify and differentiate plasmids from other circular elements.

Several software tools have been developed so far to detect and classify plasmid sequences from genomic or metagenomic data. PlasmidFinder enables precise detection and characterization of plasmid sequences based on BLAST^13^ searches against a thoroughly curated database of plasmid replicons^7^. However, this only allows detection of the replicon module, preventing the identification of other plasmid regions particularly in fragmented genomic or metagenomic assemblies. Other tools, like PlasFlow, use more complex machine learning approaches that enable the detection of plasmidic contigs based on genomic signatures of known plasmids present in current databases^14^. This approach allows identification of different plasmid regions but does not implement any classification scheme. Circular topology has been another feature used to elucidate plasmids by analyzing assembly graphs^15,16^, however, these approaches are not optimal for complex graphs like those derived from short-read metagenomic data and do not allow to differentiate from other circular elements like bacteriophage genomes. Other approaches, like those implemented by RFPlasmid^17^ and MOB-suite^18^, rely on the integration of different strategies based on database-dependent comparisons and gene conservation. More recently, the use of metagenomics to study plasmid diversity and composition directly from environmental samples, frequently referred as plasmidome analysis, has led to the development of customized pipelines that take into account Replication Initiator Proteins (RIP) domains for a more comprehensive detection of plasmids^19–21^.

Here, we introduce plaSquid, a new pipeline to identify and classify plasmid sequences from circularized or fragmented assemblies. This tool introduces a dual strategy based on: i) a database-dependent comparison algorithm optimized to differentiate chromosome from plasmid sequences with very high accuracy and, ii) an enhanced and manually-curated set of probabilistic sequence models and plasmid-specific domain architecture definitions to classify plasmids in REP types and MOB groups which outperform similar tools. Overall, we show that plaSquid allows improved plasmid detection and classification in a wide taxonomic range of bacterial genomes, diverse environmental metagenomes and targeted plasmidome sequencing analyses.

## Results

### Software overview

The plaSquid software is implemented using Nextflow^22^. Input files for plaSquid are either genomic, metagenomic or plasmidome assemblies, which can be processed by two independent but complementary approaches that are implemented as subworkflows: Repsearch and Minidist (Fig. 1). The Repsearch workflow screens assembled contigs to identify RIP domains, plasmid-specific genes and domain architectures, ctRNAs and MOB proteins based on customized and manually-curated Hidden Markov Models (HMMs) for proteins and Covariance Models (CMs) for RNA sequences. Then, filters are applied using model-specific alignment thresholds to detect and classify plasmidic contigs in REP types and mobility MOB groups (see Methods for details). The Minidist workflow aligns each assembled contig to the PLSDB plasmid database^23^ and computes a similarity index (S) to determine its plasmidic or chromosomal origin (see Methods for details).

**Figure 1.**
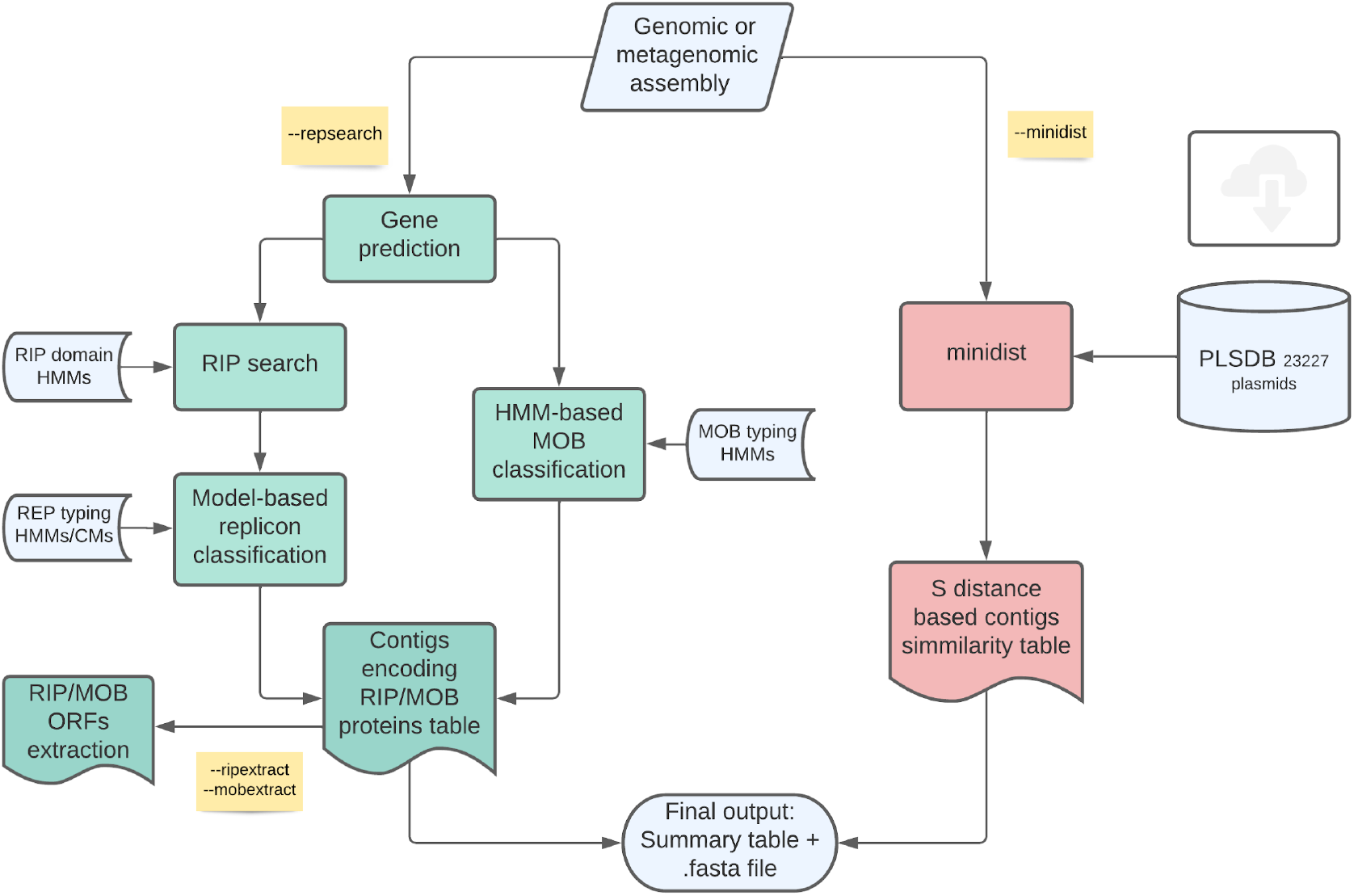
Scheme describing plaSquid pipeline. The two main workflows *repsearch* and *minidist* are shown with different colors, yellow notes describe additional options that can be passed to plaSquid Nextflow pipeline.

### Highly-accurate detection of plasmids from diverse bacterial genomes

We first aimed to compare plaSquid with current tools for plasmid detection from genomic assemblies. To do this we used a diverse genomic dataset consisting of 38 different bacterial genomes representing both Gram-negative and Gram-positive species across 10 taxonomic classes. These genomes harbor 1 to 20 plasmids (Fig. 2A; Supplementary Table S1). Then, we ran different softwares including plaSquid, PlasFlow^14^, MOB-recon^18^, PlasmidFinder^7^ and RFPlasmid^17^ to assess the precision (positive predictive value) and recall (sensitivity). Figure 2B shows that plaSquid, PlasmidFinder, MOB-recon and PlasFlow were highly-accurate (median precision > 0.9) for plasmid prediction. However, PlasFlow showed precision values as low as 0.55 for genomes belonging to the class Alphaproteobacteria. RFPlasmid exhibited the lowest precision among the tools tested (median precision = 0.63). Recall was almost optimal for plaSquid, MOB-recon and RFPlasmid which showed median values > 0.95. However, MOB-recon showed recall values < 0.5 for Alphaproteobacteria genomes. Overall, these results show that plaSquid was the only software that showed high precision and sensitivity across plasmid-containing genomes from all tested taxa.

**Figure 2.**
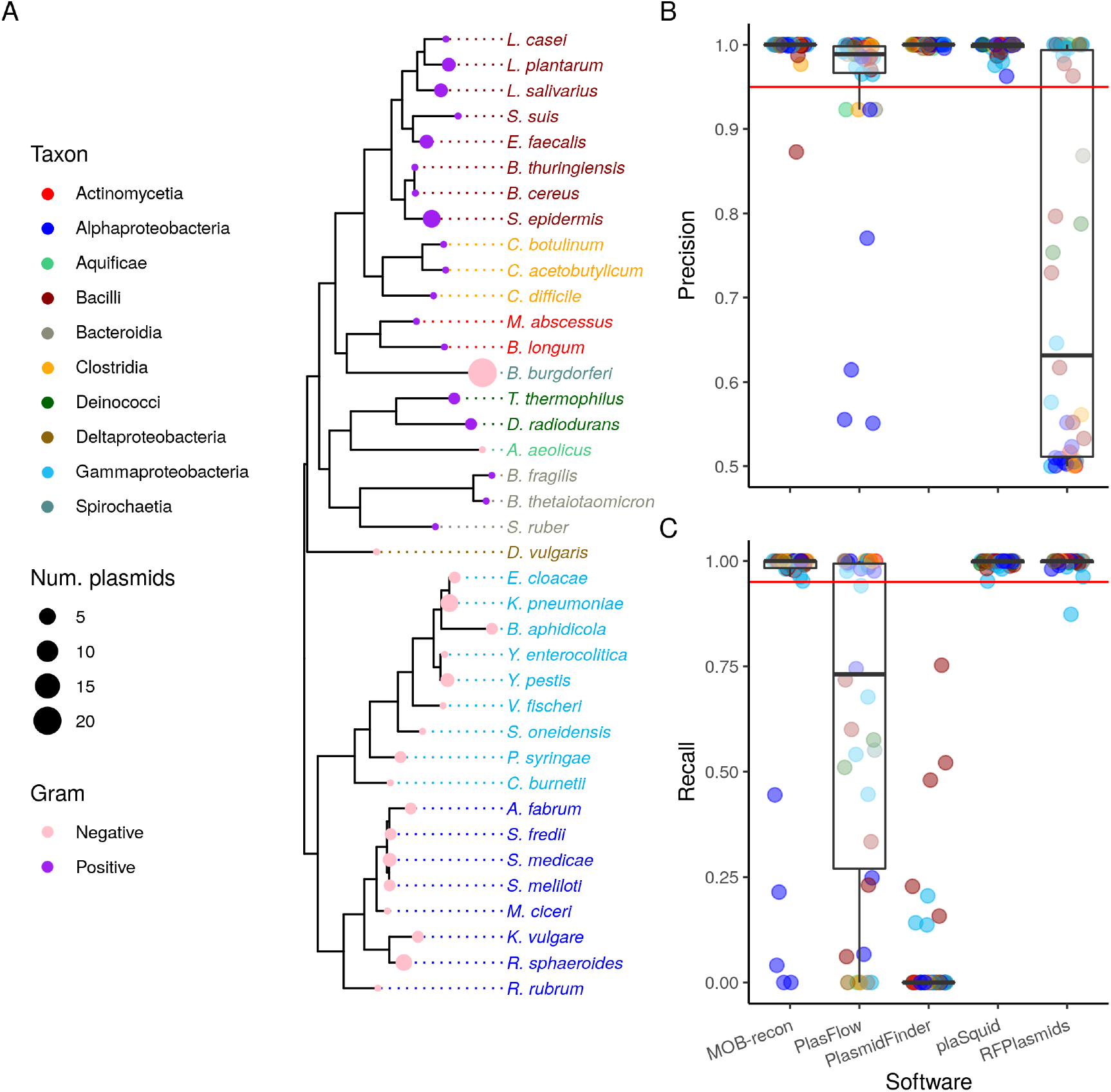
Benchmarking of plasmid detection in a diverse set of bacterial genomes. A) Phylogenetic tree based on universally conserved prokaryotic genes of the different genomes used for plasmid detection benchmarking (n = 38). Gram-negatives and Gram-positives are indicated with colored circles at tree tips, the size of the circles indicates the number of plasmids in each genome. Species names at tree tip labels are colored according to the taxonomic class of each genome analyzed. B) Boxplots showing precision values obtained for plasmid prediction with MOB-recon, PlasFlow, PlasmidFinder, plaSquid and RFPlasmid. Each point corresponds to a genome shown in the tree colored according to its taxonomic class. C) Boxplots indicating recall values obtained for plasmid predictions using the same tools and color scheme as indicated previously.

### Improved classification of replicon (REP) types

To evaluate the performance of our approach to classify REP types, we randomly sampled a manually-curated set of 469 plasmids which are representative of all known replicons (referred to as the REP reference dataset in Methods section). Then, we used this dataset to compare plaSquid results with PlasmidFinder, which is the standard approach for REP typing. As PlasmidFinder relies on BLAST^13^ searches against a predefined replicon database, we used 95%, 85% and 75% as identity and alignment coverage cutoffs. Figure 3 shows these comparisons grouping replicons according to their occurrence in a broad host range of bacteria (BHR), in Gram-positives or Gram-negatives; or according to the nature of sequences defining them (proteins or ctRNAs). Further information of grouping categories as well as correspondence between RIP domains and replicon types can be found in Supplementary Table S2.

**Figure 3.**
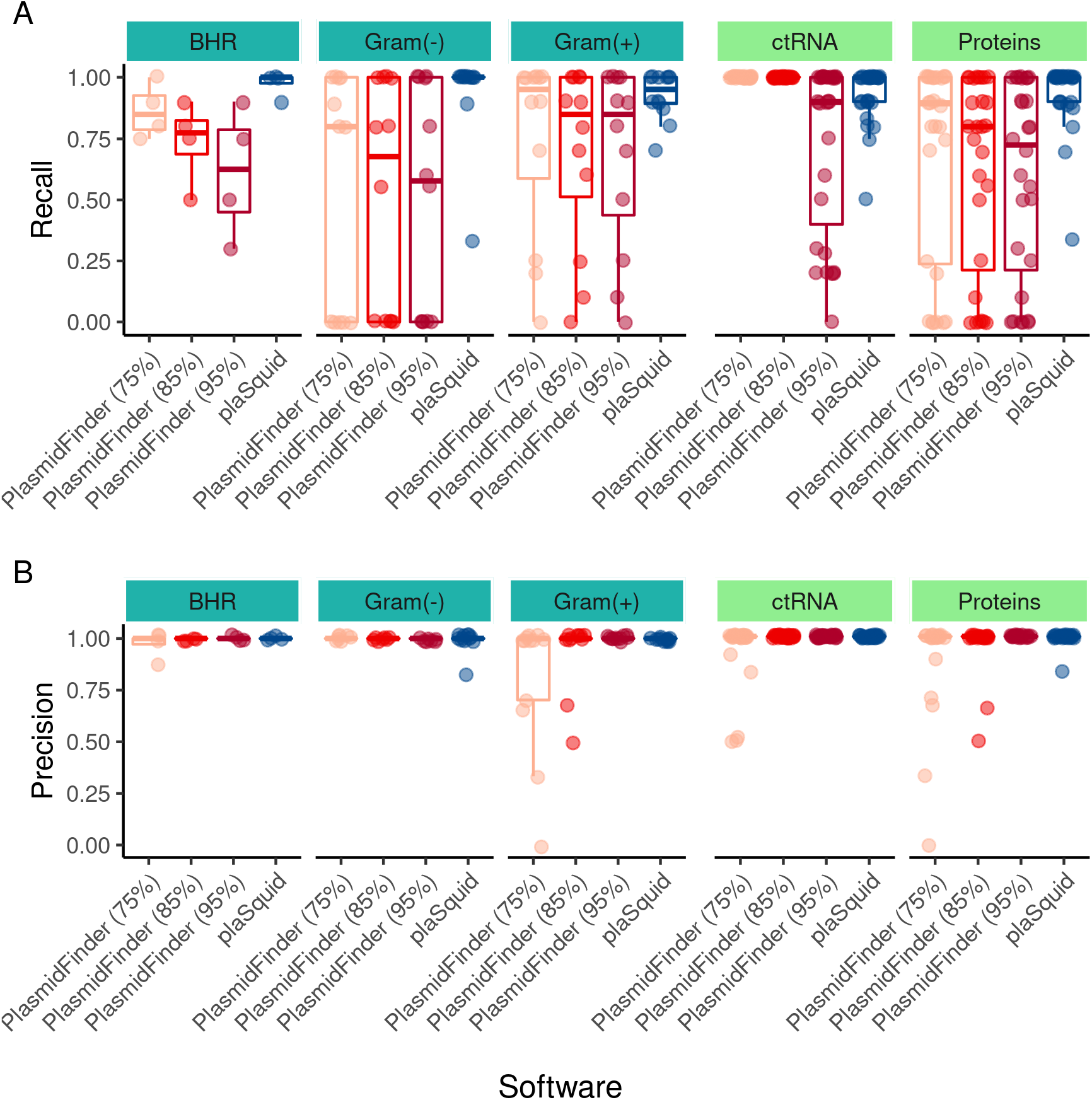
Benchmarking of replicon (REP) typing. A) Boxplots showing recall values for replicon classification benchmarking using a random sample of REP reference dataset (n = 2,065) (detailed in Supplementary Table S5). Plasmid replicons were divided according to their host taxonomy (BHR, Gram-negatives or Gram-positives) and the type of sequence determining the replicon identity (protein or ctRNA) (detailed in Supplementary Table S2). PlasmidFinder software was run with different identity and coverage thresholds (75%, 85% and 95%). B) Boxplots showing precision values for replicon classification using the same conditions as described above.

For BHR plasmids, recall values obtained with PlasmidFinder decay with more stringent cutoffs. Indeed, the best performance of this tool is observed with 75% of identity and alignment coverage (median recall = 0.8), which is lower than values obtained with plaSquid (median recall > 0.98) (Fig. 3A). For plasmid replicons exclusively found in Gram-negative bacteria, the best performance of PlasmidFinder was obtained with 75% of identity and alignment coverage (median recall = 0.74). In all tested conditions, PlasmidFinder showed greater dispersion in recall values, evidenced by a median variance of 0.12. Recall values obtained with plaSquid showed a higher median of 0.99 and a lower variance of 0.01 (Fig. 3A). For plasmid replicons found in Gram-positive bacteria, recall values were similar to the previous case. PlasmidFinder using 85% and 95% as identity and alignment coverage thresholds reported lower recall values and higher dispersion. PlasmidFinder using 75% as identity and alignment coverage thresholds and plaSquid showed comparable median recall values of 0.95 and 0.96, respectively. However, plaSquid showed a comparably lower dispersion of recall values (median variance = 0.009) (Fig. 3A).

For plasmid replicons whose classification is determined by protein sequence similarity, PlasmidFinder showed median recall values lower than 0.9 in all tested conditions, being 75% of identity and alignment coverage the best among them (median recall = 0.89). For all these conditions, PlasmidFinder showed greater dispersion of recall values (median variance = 0.16). In comparison, a substantial improvement was observed with plaSquid, evidenced by a median recall of 0.99 and a median variance of 0.02 (Fig. 3A). For plasmid replicons whose classification is determined by ctRNAs sequences, PlasmidFinder showed optimal performance when using 75% or 85% of identity and alignment coverage (median recall = 1, median variance = 0). In this case, plaSquid also showed a near-optimal median recall value of 0.99, but dispersion of values was greater than PlasmidFinder (variance = 0.01).

Figure 3B summarizes precision values obtained with plaSquid and PlasmidFinder using the same parameters as shown previously. In general, the performance of both tools in all tested conditions was near optimal, with median precision values greater than 0.99. Of note, PlasmidFinder using 75% of identity and alignment coverage showed slightly lower precision particularly for plasmid replicons found in Gram-positive bacteria (Fig. 3B).

Overall, when tested in a comprehensive set of known plasmid replicons, plaSquid outperformed the standard approach for plasmid REP typing. This improvement was particularly evident for classification sensitivity (recall).

### Improved classification of mobility (MOB) groups

To evaluate our approach for classification of MOB groups, we built a manually-curated set of 1,145 plasmids which are representative of all known MOB types (referred to as the MOB reference dataset in Methods section). Then, we used this dataset to compare plaSquid with MOB-typer. As MOB-typer relies on BLAST^13^ searches against a predefined relaxase database, we used 70%, 80% and 90% as identity and alignment coverage cutoffs. Figure 4A shows that MOB-typer using 70% as identity and alignment coverage thresholds was the condition with lowest precision (median = 0.87). The best precision with MOB-typer was obtained using 90% as identity and alignment coverage thresholds (median = 0.91). For plaSquid, precision was comparable to MOB-typer, with a median value of 0.90. Dispersion of precision values was comparable between tools, with a median variance of 4. 2 × 10^−4^ for MOB-typer (mean of the three conditions) and 4. 9 × 10^−4^ for plaSquid. Figure 4B shows recall results for the same conditions. Median recall values for MOB-typer ranged from 0.85 to 0.87, while for plaSquid this was 0.95. Dispersion of recall values was also comparable between tools, with median variance of 3. 9 × 10^−4^ for MOB-typer (mean of the three conditions) and 3. 2 × 10^−4^ for plaSquid. Together, these results evidenced that plaSquid has comparable precision but considerably better recall for classification of plasmids into MOB groups.

**Figure 4.**
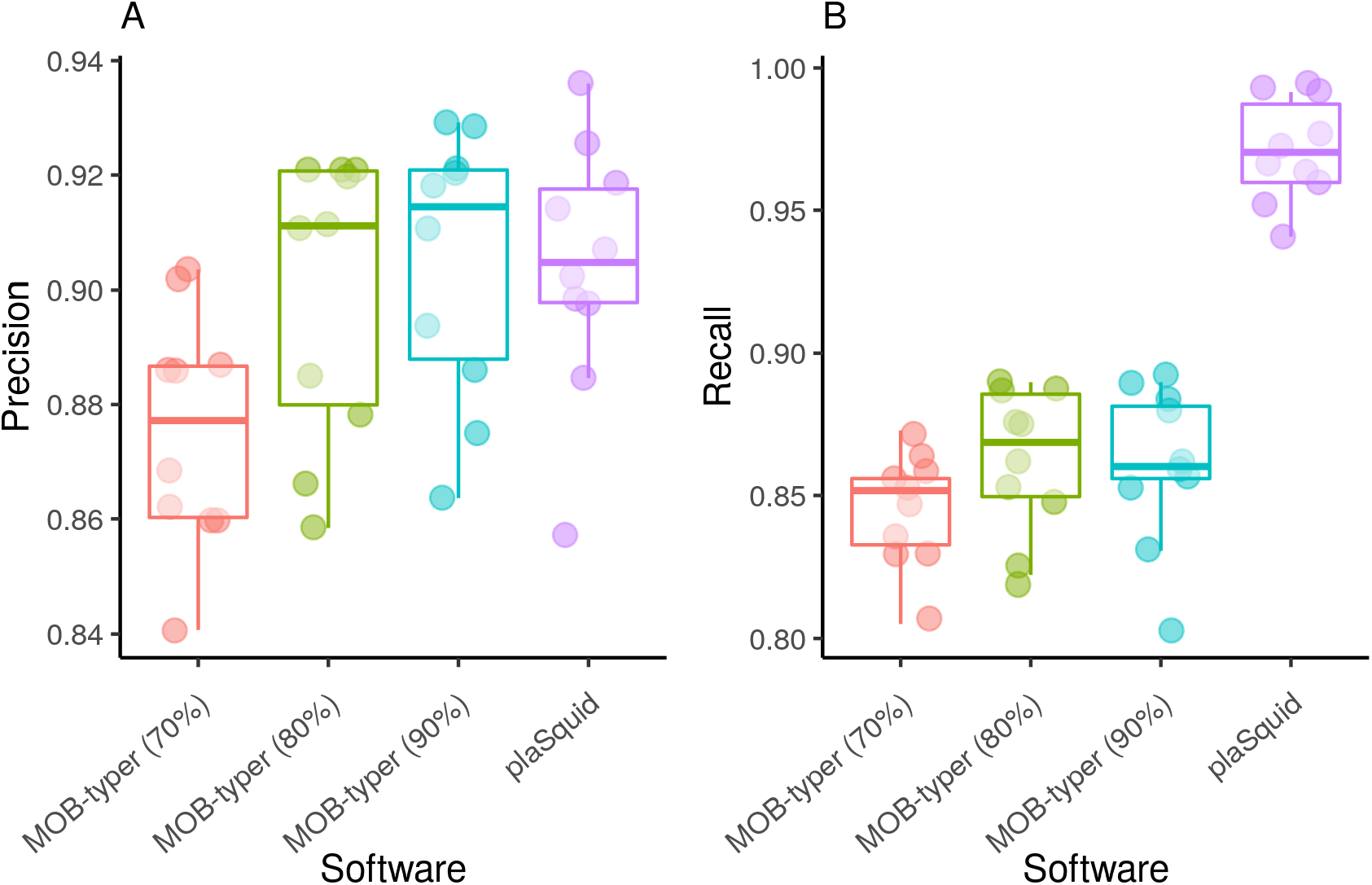
Benchmarking of mobility (MOB) groups typing. A) Boxplots showing precision values obtained with MOB-typer and plaSquid for 10 random samples of the MOB reference dataset (details in Supplementary Table S6). MOB-typer was run using 70%, 80% and 90% as sequence identity and alignment coverage thresholds. B) Boxplot showing recall values obtained with MOB-typer and plaSquid using the same conditions as above.

### Enhanced plasmid recovery from circularized plasmidome assemblies

To further assess the performance of plaSquid we used a previously published plasmidome dataset^19^. This consists of sewage water samples from 22 different countries that were processed using a plasmid-enrichment protocol and then sequenced using Oxford Nanopore. Then, the authors reported a diverse dataset of circularized elements identified as plasmids using an in-house pipeline that detects RIPs. We re-analyzed all the assembled elements (n = 165,302) using plaSquid and compared results to those reported in the original publication. Figure 5A shows the total number of plasmids detected by plaSquid (n = 78,261) is higher than the original number reported by the authors (n = 58,429). All samples analyzed with plaSquid showed a similar trend compared to the plasmid counts reported in the original publication (Fig. 5B). However, using plaSquid we detected a higher plasmid count in 19/22 (74%) of sampling locations. Overall, we observed that plaSquid detected 33.9% more assemblies as plasmids than the original report.

**Figure 5.**
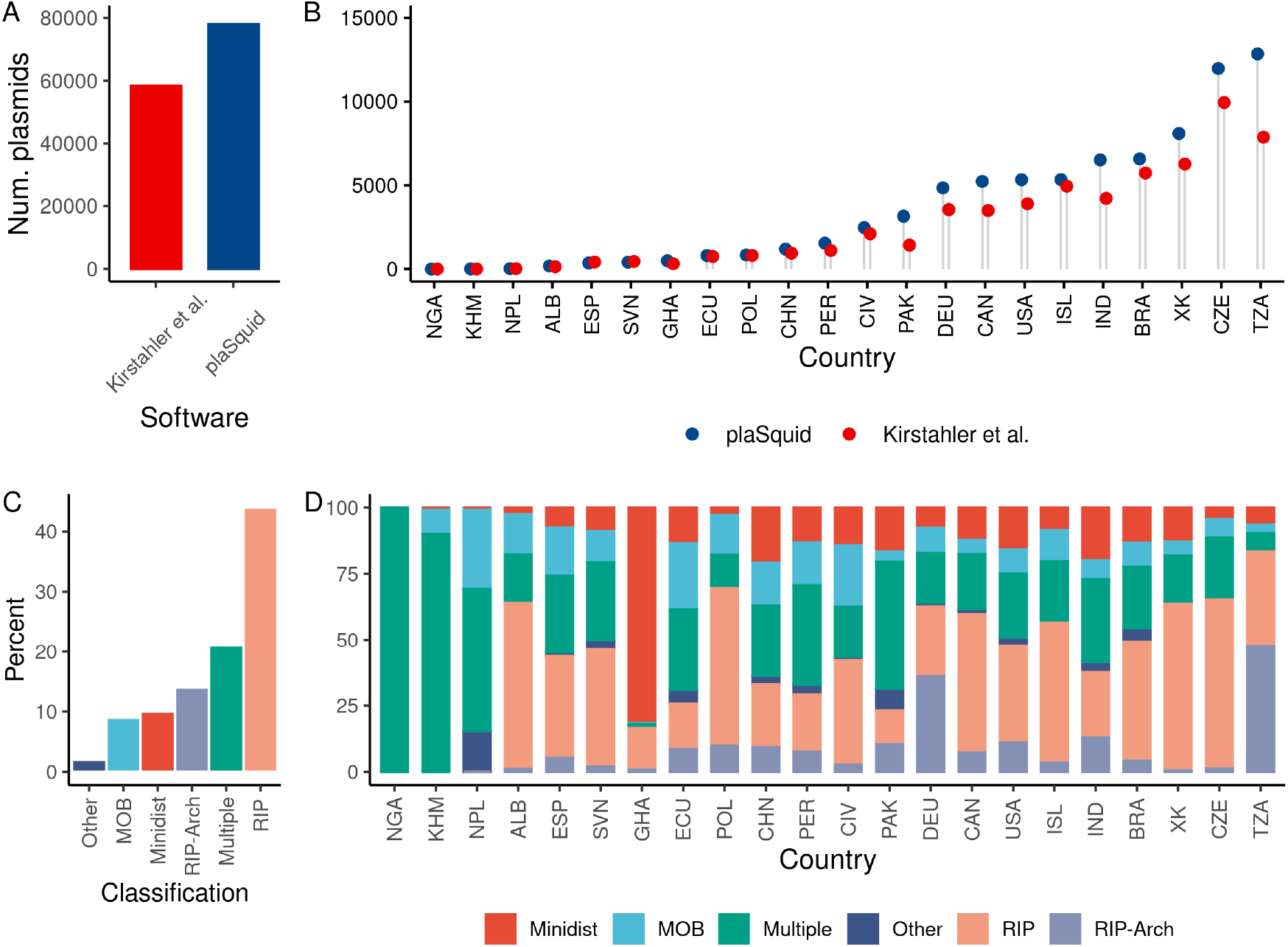
Analysis of global sewage plasmidome data using plaSquid. A) Barplot of total plasmid count recovered by plaSquid in comparison with the number originally reported by Kristahler et al. (2021)^19^. B) Plot showing the number of plasmids recovered by plaSquid in comparison to the originally reported and broken down by country. C) Barplot specifying the percentage of plasmids recovered using each different strategy implemented by plaSquid. ‘RIP’ and ‘RIP-arch’ indicate detection of single and multiple RIP domains, respectively, ‘Multiple’ indicates that a single plasmid contig was detected by more than one different detection approaches, ‘Minidist’ indicates PLSDB-dependent plasmid detection, and ‘MOB’ indicates detection of mobility groups. D) Barplot showing percentage of plasmids recovered by each strategy implemented in plaSquid broken down by country.

We also aimed to characterize the contribution of different plasmid detection strategies implemented by plaSquid using the same dataset. Figure 5C shows that the strategy based on RIP domain detection enabled the recovery of more than 40% of plasmid assemblies. Importantly, the combined contribution of Minidist and RIP-domain architectures that are new strategies incorporated by plaSquid accounted for 25% of cases. This is particularly relevant for some samples, like Brazil (BRA) in which > 90% of plasmids were identified by Minidist or Tanzania (TZN) in which > 50% of plasmids were identified by detection of RIP-domain architectures (Fig. 5D). Also, more than 20% of cases were detected using the combination of multiple strategies. Indeed, in 20 out of 22 (90%) sampling locations plaSquid identifications were based on 3 or more different approaches (Fig. 5D). Together, these results show that plaSquid enhances the recovery of plasmids from circularized, long-read assemblies generated from plasmid-enriched samples. Additionally, this improvement is explained by the integration of different detection approaches, including those newly implemented by plaSquid.

### Expanded diversity of plasmids from natural and built environments

To assess the capacity to detect new plasmid genotypes, we compared the actual diversity of plasmids found in the PLSDB database with those plaSquid was able to recover from different natural and built environments (Supplementary Table S3). To do this we measured the abundance of RIP-domain containing genes in three different metagenomic projects aiming to characterize microbes from oceans (Tara Oceans), the urban built-environment (The MetaSUB International Consortium) and the International Space Station (ISS) (Fig. 6A). From the MetaSUB dataset we were able to recover 392 Mbp of plasmid sequences, representing 42.9% of total RIP sequences found. In particular, MetaSUB was rich in RIPs containing Rep_trans, Replicase, RP-C, RP-C_C, RepL and Rep_3 domains. The Tara Oceans dataset contributed 5.3% of RIP sequences which were retrieved from 199 Mbp of plasmid sequences, with RPA and RepA_C as most prevalent domains. The ISS dataset contributed with only 1.3% of RIP sequences analyzed from 37.6 Mbp of plasmid sequences. To further confirm the observed trend, we measured the phylogenetic diversity of genes classified as each one of the different RIP domains. This analysis evidenced that plaSquid contributed to enlarging the known phylogenetic diversity in all of the 16 different RIP domains evaluated. In average, this represented an overall 2.7-fold increase in phylogenetic diversity of RIP genes (Fig. 6B). The highest expansion in phylogenetic diversity was observed for RIPs containing Rep_3, RepA_C, RP-C or RPA domains, while the lowest was observed for IncFII_repA, PriCT_1, Rep_trans, Rop, RepC and TrfA.

**Figure 6.**
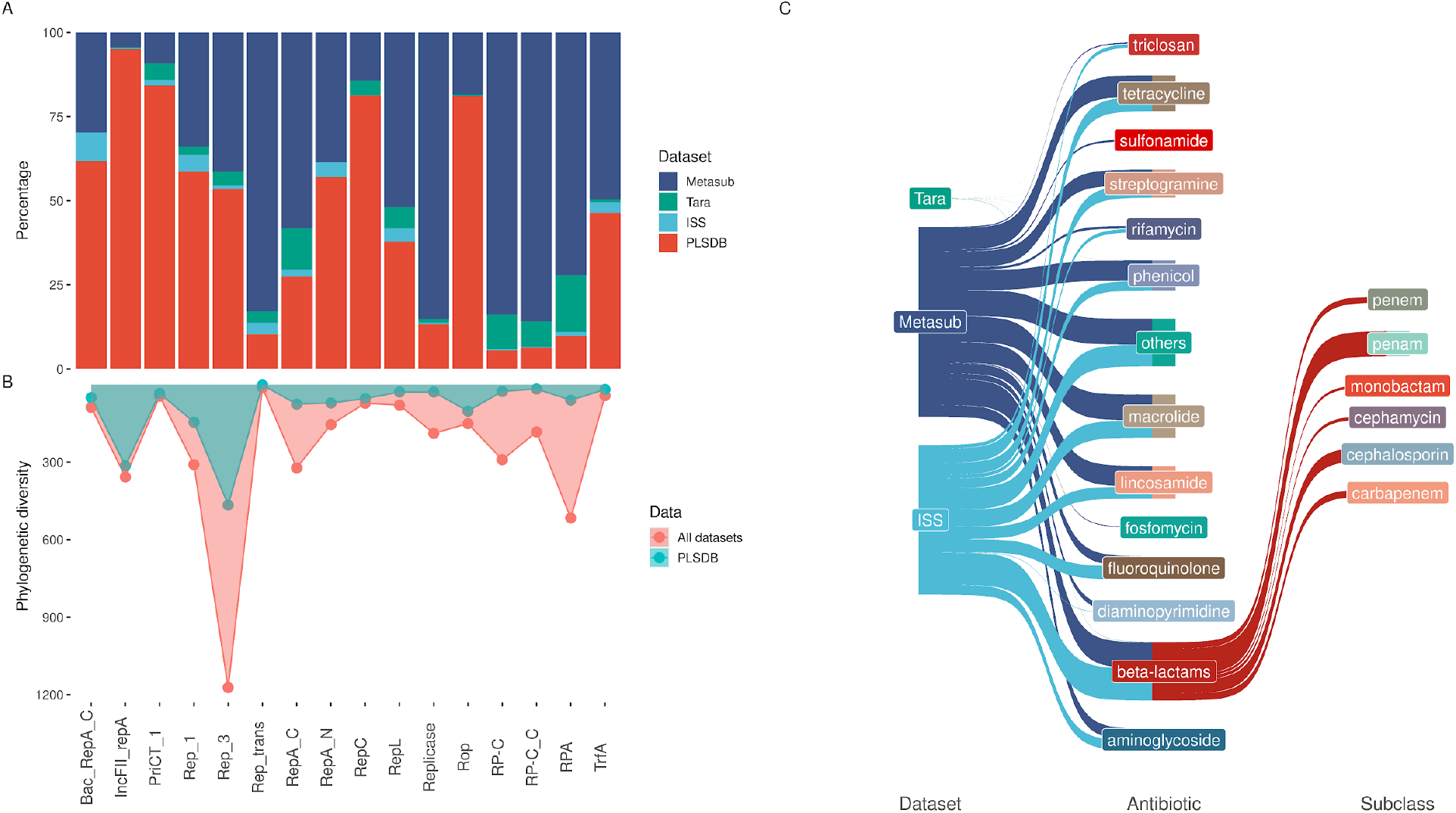
Analysis of plasmids and antimicrobial resistance applying plaSquid to diverse environmental metagenomes. A) Barplot showing the percentage of RIP sequences harboring different RIP domains in each metagenomic dataset and the PLSDB database. B) Line plot showing the plasmid diversity as measured by calculating phylogenetic diversity of each RIP type. Phylogenetic diversity was calculated for RIP sequences detected by plaSquid in the PLSDB database (pale green), and adding RIP sequences retrieved from the three metagenomic datasets analyzed (pale red). C) Sankey diagram showing antibiotic resistance found in plasmids for the three analyzed datasets. The 13 most prevailing antibiotic resistance classes are specified and beta-lactams subclasses are further shown.

Antimicrobial resistance is one of the most typical and widespread functions harbored by plasmids, so we aimed to characterize the repertoire of plasmid-derived AMR genes. Figure 6C shows ARGs found in plasmid sequences recovered from the three datasets analyzed. The Tara Oceans dataset showed very low content of ARGs in plasmid sequences, followed by the ISS and MetaSUB datasets which had similar contents. Normalization of ARG count with respect to the total number of genes in the metagenomes had an effect in favor of ISS, which had less data but still a considerable amount of ARGs in plasmid sequences. We also detected that within plasmid sequences, resistance to MLS (Macrolide, lincosamide and streptogramin), tetracycline and beta-lactams were the most enriched AMR classes (Supplementary Figure S2). Given its clinical importance, we analyzed the beta-lactam subclasses, highlighting the presence of plasmid-encoded ARGs for last-line antibiotics such as cephalosporins and carbapenem as common traits mainly in the ISS and MetaSUB datasets.

## Discussion

Here, we developed and benchmarked plaSquid, a new software tool that improves plasmid analysis by expanding detection and classification capabilities for different replicon types and hosts. Our tool leverages expert bioinformatic workflow management using Nextflow to exploit a diversified set of methods, using previously implemented^7,18,24,25^ and newly designed algorithms that enhance plasmid detection and classification from genomic or metagenomic assemblies. This approach takes advantage of the modular nature of plasmids, since different strategies can complement each other as they target distinct regions of the plasmid genome. Indeed, our benchmark analysis against four different state-of-the-art plasmid detection software confirmed that the complementary strategy implemented by plaSquid achieved the highest precision and sensitivity across a diverse set of bacterial species and plasmids.

We also introduce new methods for plasmid classification in replicon types. This approach showed better performance than PlasmidFinder, which has been the standard for classifying replicon types. We attribute this to: i) expanding the replicon repertoire, as we introduce new models for classification of certain replicon types found in plasmids from *Pseudomonas* which were not included in the PlasmidFinder database, and ii) the incorporation of specific inclusion thresholds for each classification model, enabling plaSquid to consider each replicon’s sequence diversity instead of using a fixed identity value as in PlasmidFinder. In addition, we report that changing PlasmidFinder’s similarity thresholds to values below those originally recommended by the authors helps to obtain better results with this tool. We also introduce improvements to classify plasmids in MOB groups. In this case, we integrate already generated models but added model-specific inclusion thresholds, yielding a comprehensive and precise classification of plasmid mobility which outperformed the method implemented by MOB-typer. It is worth noting that beyond we report improvements compared to standard methods, none of these classification schemes cover all known plasmid diversity. In fact, new clustering schemes based on whole-plasmid identity have recently been developed aiming to overcome this limitation^11,12,26^ and constitute complementary approaches to plaSquid.

By analyzing plasmidome data from a global sample of urban wastewater, we reinforce the notion that using multiple strategies for plasmid detection provides an overall improvement of results. Anyway, detection of RIP domains and domain architectures is the most comprehensive strategy to identify plasmids, being plaSquid the first tool that automatically incorporates this approach. Given that RIPs determine plasmid’s host range, and that previous plasmid clusters generated by whole-genome-based comparisons are mostly host-restricted^27^, we believe that the associations we uncovered between certain RIP domains and replicon types (Supplementary Table S2) could be the basis for updating and expanding the current replicon typing scheme. Hence, the important amount of information already generated for associations between replicon types and ARGs, could be better exploited to track dissemination and understand evolutionary pathways of AMR in important pathogens^28,29^.

We revealed a substantial proportion of unknown plasmids, as sequences recovered from distinct natural and built environments such as oceans, cities and the ISS greatly expanded RIPs phylogenetic diversity with respect to current plasmid databases. This is important as the rise of AMR is directly linked to human activity, and the dissemination of unknown plasmids in environments impacted by humans represent a potential reservoir for new ARGs that could emerge in the future^30^. In addition, we revealed that epidemic ARGs which today constitute important global health problems are being carried by plasmids that disseminate in our built environments. For example, resistance to betalactams had been previously reported in the ISS but was not linked to plasmid sequences^31^. Accordingly, this information may be useful to improve management of microbiological safety at the ISS and other built environments. Further exploration of this type of datasets combining plasmids and ARGs characterization could give precise information of community-wide dispersal of AMR.

Together, we consider that its scalable and reproducible implementation, the low bias in plasmid recovery from different hosts and environments given its integrative detection strategy, and the enhancements in replicon and mobility groups classification makes plaSquid suitable for large-scale and improved analysis of plasmid biology both using fragmented or complete genomic and metagenomics assemblies.

## Methods

### Replicon detection

Replicon detection is based on the identification of specific RIP protein domains from the Pfam database^32^. To build this set of RIP protein domains we reviewed recent publications that consistently used this approach to detect plasmid sequences in plasmidome studies^19,24,33^. We also added protein domains present in RIPs of specific replicon types obtained from the PlasmidFinder database^7^. Given that replication proteins similar to plasmid RIPs are also present in bacterial chromosomes, a specific inclusion threshold was set to each RIP Pfam model based on HMM bit scores obtained by comparing each model against a representative set of bacterial chromosomes from the PATRIC database^34^ and plasmid genomes from the whole PLSDB database^23^. Additionally, we used information provided by conserved domain architectures (not only the presence/absence of domains but also their linear order along the protein sequence) present in RIP proteins as a new feature to differentiate plasmid RIPs from similar replication proteins. A detailed list of domains selected for replicon detection is presented in Supplementary Table S4.

### Replicon (REP) typing

A customized and manually-curated REP reference dataset was created for replicon typing. This dataset aims to cover the diversity of different plasmid replicons that have been already classified in either a certain incompatibility group by functional analysis, or in replicon types by molecular or bioinformatic approaches. For doing this we integrated information from three different sources: i) we retrieved plasmids that had already been classified by Shintani et al. (2015)^35^, ii) we ran PlasmidFinder^7^ and MOB-typer^18^ against the PLSDB database^23^ with author’s recommended parameters, and iii) we made a thorough literature revision to identify reference sequences of different incompatibility groups that have been validated experimentally making specific emphasis in those groups currently not covered by the PlasmidFinder and MOB-typer databases. The resulting sets of sequences belonging to each replicon type were aligned using MUSCLE^36^, then alignments were manually curated and HMMs were built using HMMER v3.0^37^. Finally, we compared each sequence of each model against the same model and against the other models, aiming to define model-specific thresholds that avoid cross-detection between models. Plasmids with replicon sequences used for replicon typing are extensively detailed in Supplementary Table S5. Replicon types that are defined by RNA sequences are based on those reported by the PlasmidFinder and MOB-typer database^7^. Reference RNA sequences reported in this database were used to perform BLAST^13^ against the PLSDB database using author’s recommended identity and alignment coverage thresholds^23^. The resulting set of sequences recovered for each reference RNA defining a replicon type, was aligned using MUSCLE^36^. After that, Infernal v1.1^38^ was used to build CMs and model-specific inclusion thresholds were set by searching against the entire plasmid dataset using the same approach described for RIP models.

### Mobility (MOB) groups typing

To determine MOB groups we defined a customized and manually-curated MOB reference dataset. This consists of classifications reviewed and reported by Shintani et al. (2015)^35^, the MOB-suite reference plasmids dataset^18^ and results from searching with MOBscan tool against the PLSDB database. Together, these datasets contained plasmids belonging to MOB groups MOB_C_, MOB_F_, MOB_H_, MOB_P_, MOB_Q_ and MOB_N_. We detected that MOB_T_ and MOB_M_ groups, included in the MOBscan web tool^25^ and frequently found in integrative conjugative elements (ICEs), were also present in several plasmids of Enterococcus and Clostridioides from the PLSDB database. Accordingly, these MOB groups were also included in the final MOB reference dataset, composed of 1,145 plasmids and their associated MOB designation (Supplementary Table S6). For MOB classification, plaSquid uses HMMs reported by the MOBscan web tool, but adds model-specific classification thresholds which were set as previously explained for replicon models.

### Detection of plasmids through database comparison (Minidist)

We developed an algorithm based on sequence database comparisons to assess whether plasmid sequences can be detected and differentiated from chromosome-derived sequences. For doing this we: i) retrieved all plasmid sequences reported in the PLSDB database v2020_06_29^23^ (hereinafter referred as the plasmid database); and ii) retrieved 1,987 chromosomes from the PATRIC database that are a representative set of known bacterial taxonomic diversity^34^ (hereinafter referred as the chromosome database). To avoid spurious contamination of the chromosome database with plasmid sequences due to annotation errors, the chromosome database was manually inspected to discard sequences by looking for plasmid-specific keywords designation (i.e. “plasmid”, “replicon”, etc) (Supplementary Table S7).

From either the plasmid or chromosome databases, we took all sequences and generated 250-bp overlapping subsequences of 1.5-kb length. Then, each plasmid or chromosome 1.5-kb subsequence was mapped against the whole plasmid database using Minimap2 (-x asm5). The percentage of sequence identity and alignment coverage were used to calculate the S value that integrates the product of these two measures over the number of subsequences, according to the following equation:

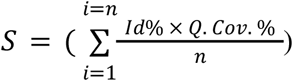

To define a value of S that maximizes the discrimination between plasmid or chromosome sequences, we took 10 random samples of 10,000 sequences each from the plasmid or chromosome database and compared S values obtained against the PLSDB database. Based on this we defined a threshold of S = 45 that optimizes classification of sequences in plasmid or chromosome origin (Supplementary Fig. S1).

### Benchmark of plasmid detection using bacterial genomes

A phylogenetically diverse dataset of plasmid-containing bacterial genomes was used to benchmark plasmid prediction in genomic assemblies (Supplementary Table S1). To avoid biases introduced by different sequencing technologies or assembly methods, complete reference genome assemblies were retrieved from the PATRIC database and were used to simulate reads with ART^39^, using default parameters. Then, simulated reads were assembled using SPAdes^40^ with default parameters and quality-checked with QUAST^41^. All resulting contigs were used as input for plasmid prediction with PlasFlow^14^, MOB-recon^18^, PlasmidFinder^7^ and RFPlasmid^17^. Plasmid prediction tools based on graph-based approaches (plasmidSPAdes^15^ and gplas^16^) were not tested because they rely on differences in sequencing depth between chromosomes and plasmids which is lost in the simulated dataset. Predictions obtained with each tool were used to calculate the proportion of reference plasmids recovered (true positives) and the proportion of reference chromosomes covered in plasmid predictions (false positives). This was used to compute precision scores for each genome and software: 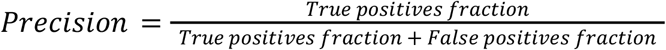. The same approach was used to calculate recall values as follows: 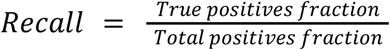.For this the total positive fraction was considered as all reference plasmids in the reference genome.

### Benchmark of replicon and MOB typing

Current tools that perform replicon typing are PlasmidFinder^7^ and MOB-recon^18^, the latter implementing a similar approach to the former. Accordingly, we compared the HMM-based approach developed for plaSquid against PlasmidFinder^7^ using 95%, 85% and 75% as identity and alignment coverage thresholds for this tool. To do this, 469 plasmids from different replicon types were equitatively sampled from the REP reference dataset in order to avoid biases in replicon type representation. The dataset contained 63 replicon categories to classify, for example IncN, while PlasmidFinder’s database contains 460 replicon-specific DNA probes, like IncN_1, IncN_2, etc. Therefore, we matched reference replicon type classifications to PlasmidFinder’s probes to compare between both methodologies (more details can be found in Supplementary Table S2). Precision and recall values were calculated as explained in the previous section. For this we considered plasmids classified in a replicon type different from the one reported in the REP reference dataset as false positives and plasmids classified in the same replicon type as in the REP reference dataset as true positives.

To assess precision and recall for mobility groups (MOB) typing, 15 plasmids of each MOB group were randomly sampled 10 times from the MOB reference dataset. These values were calculated for plaSquid and MOB-typer. For MOB-typer, different identity and alignment coverage thresholds were tested, including those recommended by the authors through personal communication (70%, 80% and 90%). False positives were determined as plasmids classified by more than one model or classified by a model not corresponding to the reference MOB group.

### Re-analysis of public metagenomic datasets

A set of 4 different metagenomic datasets generated from diverse environments and with different sequencing technologies were re-analyzed with plaSquid.

First, we retrieved plasmidome assemblies from Kristahler et al. (2021)^19^. This dataset was generated using Oxford Nanopore long-read sequencing after DNA extraction and plasmid enrichment from sewage samples of 22 different countries across the world. Resulting sequences have been processed and assembled by the authors aiming to generate circular contigs. Then, resulting circular contigs were screened for the presence of plasmid-related genes to confirm true plasmid sequences using an in-house pipeline reported in the original paper. All circular contigs reported by the authors were used as input to plaSquid to determine their plasmidic origin. We also investigated the contribution of each strategy implemented in plaSquid for plasmid detection.

Second, we analyzed 3 different sets of shotgun metagenomic assemblies generated from short reads sequencing technologies. These samples were collected from diverse environments: i) the Tara Oceans Expedition dataset consisting of secondary assemblies from 264 samples sites representing 10 oceanographic provinces across the world^42^; ii) a subset of the public transport surface metagenomic dataset from the MetaSUB International Consortium^43^, representing samples from 51 cities around the world; and iii) 50 metagenomes generated from surface and air filter samples collected in the International Space Station (ISS)^31^. All contigs reported in these datasets were filtered to a minimum of 1500 base pairs and used as input to plaSquid in order to detect and classify plasmid contigs. Plasmids contained in the PLSDB database were also analyzed with plaSquid in order to compare against these datasets. Additionally, we used the *--ripextract* sub workflow which enables the automatic extraction of RIP domain-containing ORFs for further analysis of these proteins. Resulting RIP sequences from the analysis of PLSDB and the three metagenomic datasets were clustered at 90% of identity with CD-HIT^44^ for each domain. Representative sequences of each RIP-domain cluster were aligned using the msa package^45^, phylogenetic trees were computed with the neighbor-joining algorithm of ape package^46^ and visualized using the ggtree package^47^. The sum of branch lengths (Faith’s phylogenetic diversity) was computed for each tree of RIP sequences found for each domain.

Antibiotic resistance genes were detected using ABRicate software (https://github.com/tseemann/abricate) with the CARD database^48^ using 80% identity and coverage as inclusion thresholds. The tidyverse package^49^ was used to develop a custom script to extract ARGs of plasmid origin. The fraction of ARGs genes in plasmids was calculated by dividing by the total number of genes annotated in plasmid contigs.

## Acknowledgements

This work was partially supported by the G4 program from Institut Pasteur de Montevideo funded by the Banco de Seguros del Estado (BSE) of Uruguay. M.G. and I.F. are supported by the Agencia Nacional de Investigación e Innovación of Uruguay (ANII), grant numbers POS_FSA_2019_1_1008860 and POS_NAC_2018_1_151494, respectively.

## Author Contributions

G.I. and M.G. conceived the idea. M.G. wrote the software with inputs from I.F. M.G. performed the analyses with inputs from G.I. M.G. prepared the first version of the manuscript. M.G., I.F. and G.I. wrote and approved the final version of the manuscript.

## Competing Interests

Nothing to declare.

## Data Availability

Source code is available at plaSquid Github repository https://github.com/mgimenez720/plaSquid under the GPLv3 license. Reference chromosome dataset is available at https://figshare.com/s/507c91ae930e9740cbde. The REP reference dataset is available at https://figshare.com/s/07747fafed60e59f1992. The MOB reference dataset is available at https://figshare.com/s/12f51c5433b9049a8b43. Other supplementary files and tables are available upon request.

## Notes

https://github.com/mgimenez720/plaSquid

### Competing Interest Statement

The authors have declared no competing interest.

